# Rapid visual CRISPR assay: a naked-eye colorimetric detection method for nucleic acids based on CRISPR/Cas12a and convolutional neural network

**DOI:** 10.1101/2021.07.17.452802

**Authors:** Shengsong Xie, Dagang Tao, Yuhua Fu, Bingrong Xu, You Tang, Lucilla Steinaa, Johanneke D Hemmink, Wenya Pan, Xin Huang, Xiongwei Nie, Changzhi Zhao, Jinxue Ruan, Yi Zhang, Jianlin Han, Liangliang Fu, Yunlong Ma, Xinyun Li, Xiaolei Liu, Shuhong Zhao

## Abstract

Rapid diagnosis based on naked-eye colorimetric detection remains challenging, but it could build new capacities for molecular point-of-care testing (POCT). In this study, we evaluated the performance of 16 types of single-stranded DNA-fluorophore-quencher (ssDNA-FQ) reporters for use with CRISPR/Cas12a based visual colorimetric assays. Among them, 9 ssDNA-FQ reporters were found to be suitable for direct visual colorimetric detection, with especially very strong performance using ROX-labeled reporters. We optimized the reaction concentrations of these ssDNA-FQ reporters for naked-eye read-out of assay results (no transducing component required for visualization). Subsequently, we developed a convolutional neural network algorithm standardize and to automate the analytical colorimetric assessment of images and integrated this into the MagicEye mobile phone software. A field-deployable assay platform named RApid VIsual CRISPR (RAVI-CRISPR) based on a ROX-labeled reporter with isothermal amplification and CRISPR/Cas12a targeting was established. We deployed RAVI-CRISPR in a single tube towards an instrument-less colorimetric POCT format that requires only a portable rechargeable hand warmer for incubation. The RAVI-CRISPR was successfully used for the single-copy detection of severe acute respiratory syndrome coronavirus 2 (SARS-CoV-2) and African swine fever virus (ASFV). Our study demonstrates this novel RAVI-CRISPR system for distinguishing different pathogenic nucleic acid targets with high specificity and sensitivity as the simplest-to-date platform for rapid pen-side testing.

## Introduction

Nucleic acid-based assays have been widely adopted in many fields, for example, in the diagnosis of infectious diseases^1^, genotyping^2^, and food safety testing^3^. Infectious diseases are presently the focus of considerable research attention to resolve long-standing public health problems from "classical" diseases to newly emerging epidemic outbreaks. In particular, severe acute respiratory syndrome coronavirus 2 (SARS-CoV-2), the novel RNA virus responsible for the COVID-19 pandemic, has a major impact on human health worldwide^4^, infecting over 178 million people as of June 2021, reported to the World Health Organization (WHO) (https://covid19.who.int). In order to control COVID-19 pandemic, there is an urgent need to test for the SARS-CoV-2 in as many people as possible^5,6,7^. Similarly, animal production is periodically subject to widespread viral epidemics, as in the case of African swine fever virus (ASFV), a DNA virus that infects domestic pigs and wild boars with a mortality rate of up to 100%^8^. Although agricultural epidemics receive less attention than human epidemics, the early diagnosis of ASFV is necessary to monitor circulating virus for the control and prevention of current and future outbreaks^9,10^, which have major impacts on food production and economy^11^. Although current PCR-based clinical diagnostic methods for the detection of viral nucleic acids are highly sensitive, these methods require expensive equipment, trained technicians, and clean laboratory environments, thus limiting their application in developing nations and rural areas without clinical infrastructure^12^.

Thus, increasing the affordability of point of care testing (POCT)^13^, and moving diagnostic testing for viral pathogens such as COVID-19 or ASF from the laboratory to pen-side detection could be potentially transformative for epidemiological efforts. Although real-time quantitative PCR (qPCR)-based nucleic acid detection is the current gold standard for SARS-CoV-2 and ASFV diagnosis^14,15^, isothermal amplification techniques, such as Loop-Mediated Isothermal Amplification (LAMP) and Recombinase Polymerase Amplification (RPA), represent comparatively easy, accessible, and reliable alternatives^16,17^. In particular, these methods can combine a variety of portable read-out types for POCT, such as paper lateral flow test strips or fluorescence and colorimetric assays^18,19,20^. LAMP/RPA-based detection methods have been rapidly developed and deployed for both SARS-CoV-2 or ASFV owing to their high efficiency of amplification^21,22,23,24^. However, the vast majority of these LAMP/RPA-based assays depend on non-specific nucleic acid detection, meaning that unintended LAMP/RPA amplicons cannot be distinguished from the amplicons of correct targets, leading to unreliable test results^25,26^.

To overcome this issue, nucleic acid detection methods based on combination of Clusters of Regularly Spaced Short Palindrome Repeats (CRISPR) and LAMP or RPA were developed, including SHERLOCK, DETECTR, HOLMES, and CDetection^27,28,29,30^. These diagnostic strategies use Cas proteins, which have collateral nuclease activity to specifically target double-strand DNA (dsDNA) or single-strand RNA (ssRNA) and a protospacer adjacent motif (PAM) sequence, thereby effectively avoiding false positive results in isothermal amplification^31,32^.

Moreover, some studies have reported that Cas-based detection methods can distinguish differences at the single nucleotide level by integrating an optimized CRISPR RNA (crRNA)^29,30,33^, and several improvements to CRISPR-based detection have been developed for POCT applications, such as CRISPR-based lateral flow strip assay^34,35^, CRISPR-based fluorescent cleavage assay^36,37^, plasmonic CRISPR/Cas12a assay^38,39^, and Cas12a-modulated fluorescence resonance energy transfer assay that incorporates nanomaterials for nucleic acid sensing^40^. In addition, mobile phone applications (apps) and mobile phone-based devices have been developed to support the evaluation of lateral flow strips or fluorescence signal read-outs^41,42,43^. Other studies have established methods that seek to simplify the reaction process like the All-In-One Dual CRISPR/Cas12a assay^44^ and a CRISPR/Cas12a based portable biosensor^45^.

In this study, to establish an accurate and convenient method based on colorimetric detection of viral nucleic acids, we developed a practical POCT detection system that does not require advanced training or specialized instruments. We demonstrated the proof-of-concept in RApid VIsual CRISPR (RAVI-CRISPR) system for sensitive testing of simulated SARS-CoV-2 and real ASFV infection samples by a naked-eye detection. Subsequently, we developed a convolutional neural network algorithm to standardize and automate the analytical colorimetric assessment of images and integrated this into a MagicEye mobile phone software for rapid detection of SARS-CoV-2 and ASFV.

## Results

### Establishment of the RAVI-CRISPR assay

To identify the best reporter for the RAVI-CRISPR assay, we screened a total of 16 types of ssDNA-FQ reporters using an exonuclease I digestion and subsequently optimized their concentrations for direct application in RAVI-CRISPR assays (Supplementary Table S1). The two main steps in the assay consisted of an exonuclease I digestion and a reporter signal read-out (Fig. 1A). The colors emitted by different types of degraded ssDNA-FQ reporters in reaction solution were then compared under a blue or UV light transilluminator, as well as with direct assessment by the naked eye using no excitation light. In addition, the fluorescence signal intensities of the intact or degraded ssDNA-FQ reporters were measured using a fluorescence microplate reader to establish background signals. The results showed that 14 of the degraded ssDNA-FQ reporters can emit strong fluorescence signals under the UV light transilluminator (Fig. 1B).

**Figure 1.**
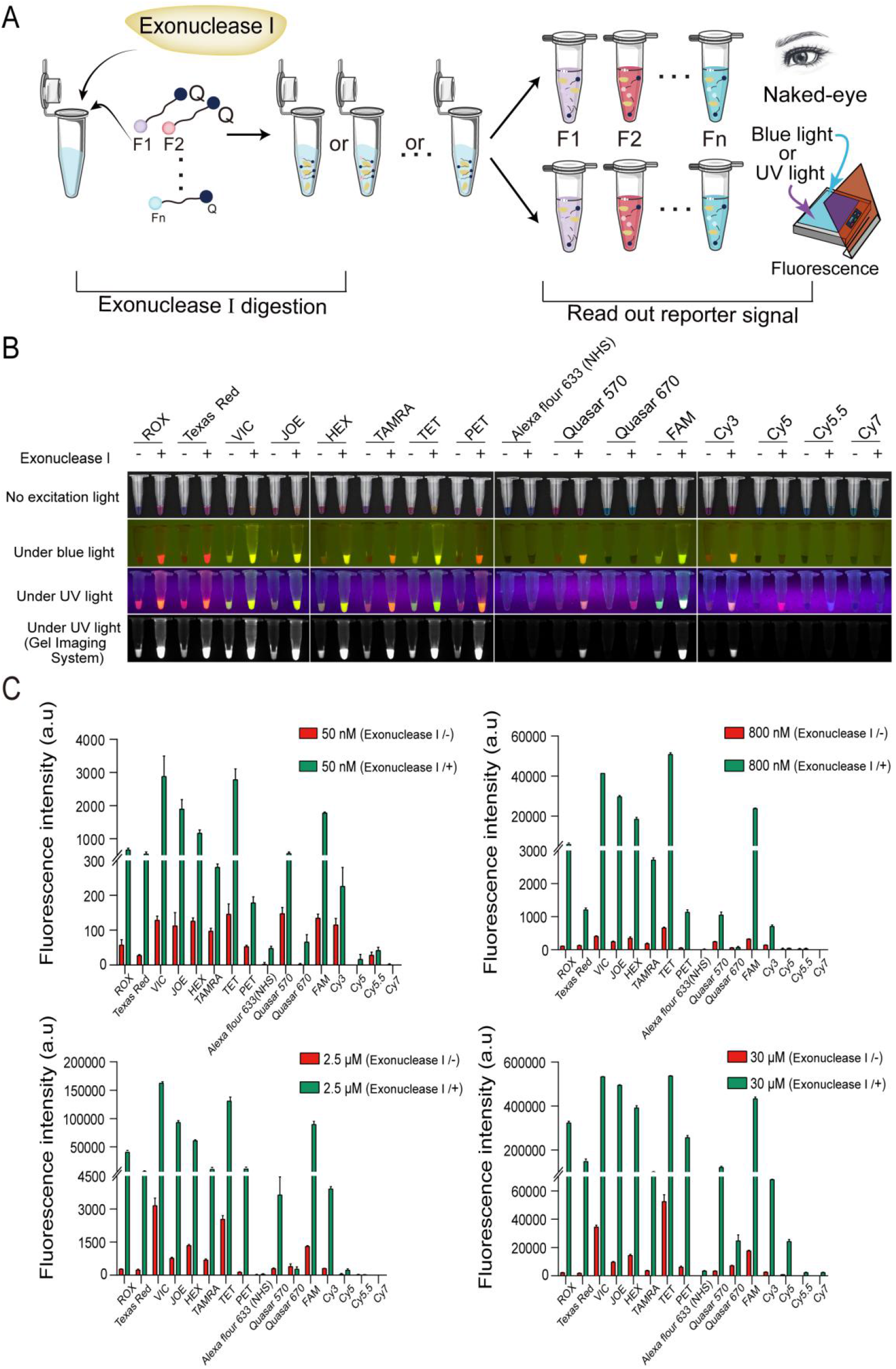
Screening and optimization of the ssDNA-FQ reporters for RAVI-CRISPR assays. **A.** Schematic diagram of the assay for determining ssDNA-FQ reporter activity based on exonuclease I cleavage (left) and subsequent evaluation under the blue or UV light or by the naked eye observation of colorimetric changes in the reaction solution (right). **B.** Visual and end point imaging evaluation of the excitation or colorimetric characteristics of 16 candidate ssDNA-FQ reporters after exonuclease I cleavage, 15 min incubation, and thermal denaturation. + represents a reaction with exonuclease I, - represents a reaction without exonuclease I. **C.** Optimization of ssDNA-FQ reporter concentrations in RAVI-CRISPR assays. A fluorescence microplate reader was used to quantify the fluorescence intensity and background signals for individual ssDNA-FQ reporters at four concentrations (50 nM, 800 nM, 2.5 μM, and 30 μM), both before and after exonuclease I cleavage. Data are represented as means ± SEM; n = 3. The images were captured under the blue (470 nM) or UV lights by a smartphone camera or the gel imaging system. The images were also captured under no excitation light by a smartphone camera.

Interestingly, we observed by the naked eye that more than 9 of the degraded ssDNA-FQ reporters changed the color of the reaction solution in the PCR test tube without excitation light (Fig. 1B, Supplementary Table S1). For instance, the degraded ROX- or Texas RED-labeled reporter reactions exhibited obvious color changes from blue to red (Fig. 1B). However, the degraded Cy5- and Cy5.5-labeled reporters were only found to emit fluorescence signals under UV light, while the degraded Cy7-labeled reporter did not emit any fluorescent signal nor change the color in reaction solutions (Fig. 1B). In conclusion, these results indicated that some ssDNA-FQ reporter modifications had better optical properties than others.

In the naked eye detection of nucleic acids, the optimal concentrations of ssDNA-FQ reporters significantly improved the visual colorimetry (Supplementary Fig. S1A-1P). However, for enhanced fluorescence visualization assays, background noise was concurrently increased with increasing signals for VIC, TET, and FAM-labeled reporters, resulting in potential interference with detection accuracy (Supplementary Fig. S1A-1P). We further confirmed, using a fluorescence microplate reader, that for some VIC, TET or FAM-labeled ssDNA-FQ reporters, higher concentrations resulted in excessive background fluorescence signals (Fig. 1C). In conclusion, an optimal concentration should be determined and adjusted for each ssDNA-FQ reporter type depending on the method used to assess the read-out by either fluorescence excitation or the naked eye evaluation with no excitation light. For instance, for a more accurate colorimetric naked eye detection, the optimal concentration was the ROX-labeled reporter at ≥ 5 μM (Supplementary Fig. S1A). However, for the detection of fluorescence intensity under blue light transilluminator, the minimal concentration of ROX-labeled reporter was 100 nM after 15 min incubation to obtain a full saturation (Supplementary Fig. S1A). Given the strong performance of the ROX dye, we selected the ROX-labeled reporters for all subsequent experiments.

### SARS-CoV-2 detection by the RAVI-CRISPR assay

In order to develop a highly sensitive diagnostics for the nucleic acids of SARS-CoV-2 using the RAVI-CRISPR system and the naked eye detection, five crRNAs for the SARS-CoV-2 *E* gene (E-crRNA-1 to 5) and nine crRNAs for the *N* gene of SARS-CoV-2 (N-crRNA-1 to 9) were screened for their efficiency of target cleavage (Supplementary Fig. S2A). First, we compared the activities of these crRNAs using PCR products and corresponding ssDNA activators as templates. The results showed that E-crRNA-2 and E-crRNA-5 exhibited the highest activity for detection of the *E* gene (Supplementary Fig. S2B), while N-crRNA-5, N-crRNA-6, N-crRNA-7, and N-crRNA-9 showed sufficient activity for detection of the SARS-CoV-2 *N* gene (Supplementary Fig. S2C). Subsequent experiments used only E-crRNA-5 and N-crRNA-9 for the detection of SARS-CoV-2.

To further evaluate the sensitivity of the RAVI-CRISPR technology for SARS-CoV-2 detection, *in vitro* transcribed RNAs of the *N* and *E* genes from two synthetic plasmids, each carrying one of the target genes, were used as a template (Fig. 2A). Following a concentration gradient, we determined that the limit of detection (LoD) for amplification by the RT-LAMP assay was 40 copies/μL (Fig. 2B). Subsequently, the amplified product of RT-LAMP *N* gene was digested by Cas12a and subjected to the detection. The result showed that the sensitivity of this technique reached 40 copies/μL (Fig. 2C). The fluorescence signal intensity of the RAVI-CRISPR assay was further measured using a fluorescence microplate reader (Fig. 2D), which confirmed the detection by the naked eye with or without excitation light. In addition, we tested both a lateral-flow paper strip assay integrated with CRISPR/Cas12a (Fig. 2E) as well as a traditional quantitative reverse transcription PCR (RT-qPCR) assay (Fig. 2F), the results of both assays were consistent with the sensitivity of the direct naked eye detection without excitation light. For example, we observed that RT-qPCR Ct values were ≥ 40 for sample concentrations ≤ 4 copies/μL (Fig. 2F, Supplementary Table S2), which can be considered as negative results. Furthermore, we found that the RAVI-CRISPR technology can directly detect the *N* gene of SARS-CoV-2 by the naked eye assessment but cannot detect its orthologue *N* gene of SARS-CoV or MERS-CoV (Fig. 2G). This specificity was consistent with the SARS-CoV-2 *N* gene detected by CRISPR/Cas12a combined with a lateral-flow paper strip (Fig. 2H).

**Figure 2.**
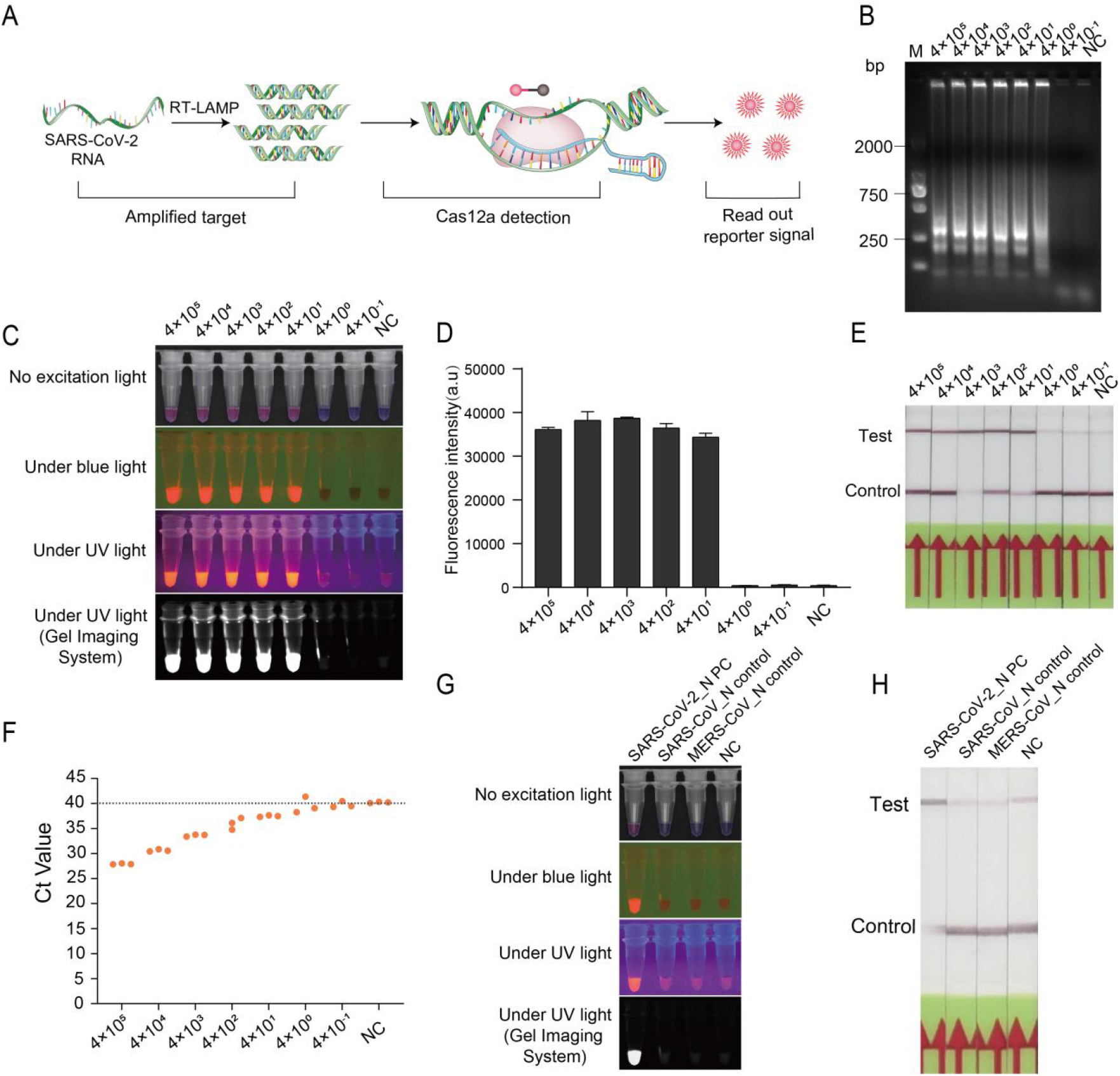
The sensitivity and specificity of RAVI-CRISPR assay for the detection of *in vitro* SARS-CoV-2 *N* gene transcripts. **A.** Schematic diagram of the RAVI-CRISPR assay for the detection of SARS-CoV-2. **B.** Agarose gel electrophoresis determination of the limit of detection for RT-LAMP amplification of the SARS-CoV-2 *N* gene. **C.** Colorimetric signal detection of a 10-fold serial dilution of *in vitro* SARS-CoV-2 *N* gene transcripts using the RAVI-CRISPR assay. **D.** Sensitivity of the RAVI-CRISPR assay quantified with a multifunctional microplate reader. **E.** Sensitivity of the RAVI-CRISPR-based lateral flow strip assay. **F.** Sensitivity of the detection of *in vitro* SARS-CoV-2 *N* gene transcripts by RT-qPCR using an CFX96 Touch Real-Time PCR Detection System. **G.** Specificity of the RAVI-CRISPR assay evaluated by the naked eye or fluorescent visual detection. **H.** Specificity of the RAVI-CRISPR-based lateral flow strip assay. Data are represented as means ± SEM; n = 3. NC stands for negative control while PC stands for positive control.

We next examined the sensitivity and specificity of SARS-CoV-2 *E* gene detection by the RAVI-CRISPR assay. Using the concentration gradient, we determined the LoD for RT-LAMP amplification of the SARS-CoV-2 *E* gene to be 58 copies/μL (Supplementary Fig. S3A). The amplified product of the *E* gene was then digested by Cas12a for the naked eye detection under different light conditions. The result showed that the sensitivity of this technique reached 58 copies/μL (Supplementary Fig. S3B), which was confirmed by microplate reading (Supplementary Fig. S3C). As with the *N* gene, both CRISPR/Cas12a combined with lateral-flow paper strip (Supplementary Fig. S3D) and traditional RT-qPCR assays (Supplementary Fig. S3E, Table S2) showed comparable sensitivity for the detection of SARS-CoV-2 *E* gene with that of the RAVI-CRISPR assay. Specifically, the Ct values observed in the RT-qPCR assay at the sample concentration ≤ 5.8 copies/μL were ≥ 40 (Supplementary Fig. S3E), indicating a negative result. These results confirmed the sensitivity of this technique to be 58 copies/μL for the SARS-CoV-2 *E* gene. Finally, we found that the RAVI-CRISPR system has high specificity for detecting the SARS-CoV-2 *E* gene and it can clearly distinguish the targeted *E* gene from different SARS family viruses (Supplementary Fig. S3F). This result was further confirmed with the CRISPR/Cas12a - lateral flow paper strip combination assay (Supplementary Fig. S3G). Thus, the RAVI-CRISPR systems using RT-LAMP coupled with CRISPR/Cas12a allowed a robust detection of the SARS-CoV-2 *E* and *N* genes with high sensitivity and specificity.

### ASFV detection by the RAVI-CRISPR assay

Considering our result showing effective detection of SARS-CoV-2, we next developed a fast, sensitive, and reliable RAVI-CRISPR-based assay for ASFV using a highly active crRNA identified in our previous study37. The strategy for ASFV detection was similar to that for SARS-CoV-2 except that a standard LAMP for DNA amplification was used rather than RT-LAMP since ASFV is a DNA virus (Fig. 3A). We first evaluated the sensitivity of ASFV *p72* by RAVI-CRISPR visual detection using a concentration gradient to determine the LoD to be 7 copies/μL (Fig. 3B). Subsequently, we assessed our system using 21 clinical samples previously diagnosed using a qPCR recommended by the World Organisation for Animal Health (OIE). The result showed 100% consistency between the naked eye assessment of RAVI-CRISPR-based assay under different light conditions (Fig. 3C) and the qPCR result (Fig. 3D) for both negative and positive samples. Notably, the negative samples with Ct values observed in the OIE qPCR ≥ 35 were confirmed by the RAVI-CRISPR assay (Supplementary Table S3).

**Figure 3.**
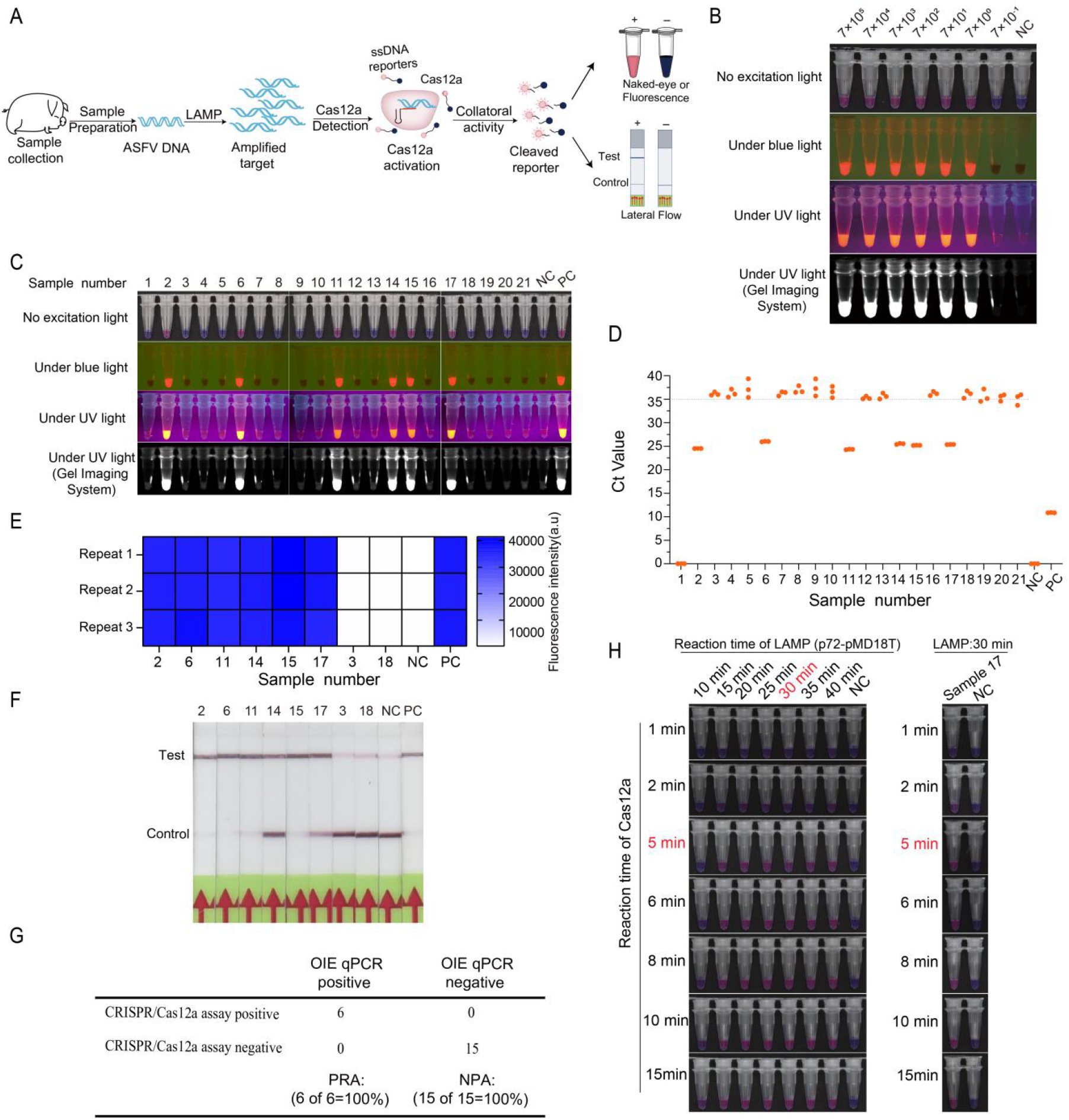
The sensitivity and specificity of RAVI-CRISPR assay for the detection of ASFV positive samples. **A.** Schematic diagram of steps in the RAVI-CRISPR assay for the detection of ASFV. **B.** The sensitivity of the RAVI-CRISPR assay determined using 10-fold serial dilutions of ASFV positive samples. **C.** The naked eye detection of ASFV positive samples by the RAVI-CRISPR assay under blue, UV or no excitation light conditions. **D.** Detection of ASFV positive samples by qPCR (n = 21) using an CFX96 Touch Real-Time PCR Detection System. **E.** Microplate reader quantification of selected ASFV positive samples confirmed by the RAVI-CRISPR assay. **F.** RAVI-CRISPR-based lateral flow strip assay detection of selected ASFV positive samples confirmed by the RAVI-CRISPR assay. **G.** Comparison of the detection results of selected ASFV positive samples by the RAVI-CRISPR-based and OIE qPCR-based assays. PPA stands for positive predictive agreement while NPA stands for negative predictive agreement. **H.** Evaluation of different reaction incubation times for LAMP amplification and CRISPR cleavage for the RAVI-CRISPR assay. NC stands for negative control while PC stands for positive control.

Moreover, the read-outs of the RAVI-CRISPR assay were further validated by a multifunctional microplate reader (Fig. 3E) and the CRISPR/Cas12a-integrated lateral-flow paper strip assay (Fig. 3F). All these results indicated that the detection of ASFV *p72* using the CRISPR/Cas12a-based assay can be accurately applied in the clinical screening of ASF samples (Fig. 3G). In addition, we evaluated the optimal reaction time for RAVI-CRISPR read-out using a p72-containing plasmid as template. The result showed that the shortest reaction time necessary for the LAMP reaction to reach saturation was around 30 min, while the minimum reaction time required for Cas12a cleavage in the naked eye detection was around 5 min (Fig. 3H). Therefore, this optimized RAVI-CRISPR assay including only 35 min of incubation can detect as few as 7 copies/μL of virus particles per reaction, which was further verified by the screening of ASF infection samples (Fig. 3H).

### Convolutional neural network assessment of the read-out of RAVI-CRISPR assay

In order to provide the RAVI-CRISPR assay as a convenient package including a standardized evaluation mode, we also developed the MagicEye mobile phone app that employed two machine learning models based on convolutional neural networks to automatically interpret the results of nucleic acids detection in multiple reaction tubes (Fig. 4). The first model was for object detection, which used a Single Shot MultiBox Detector (SSD) algorithm to identify the images of reaction tubes^46^. The second model was for binary classification of either positive or negative signals to resolve the operator-associated inconsistencies across the naked eye detection.

**Figure 4.**
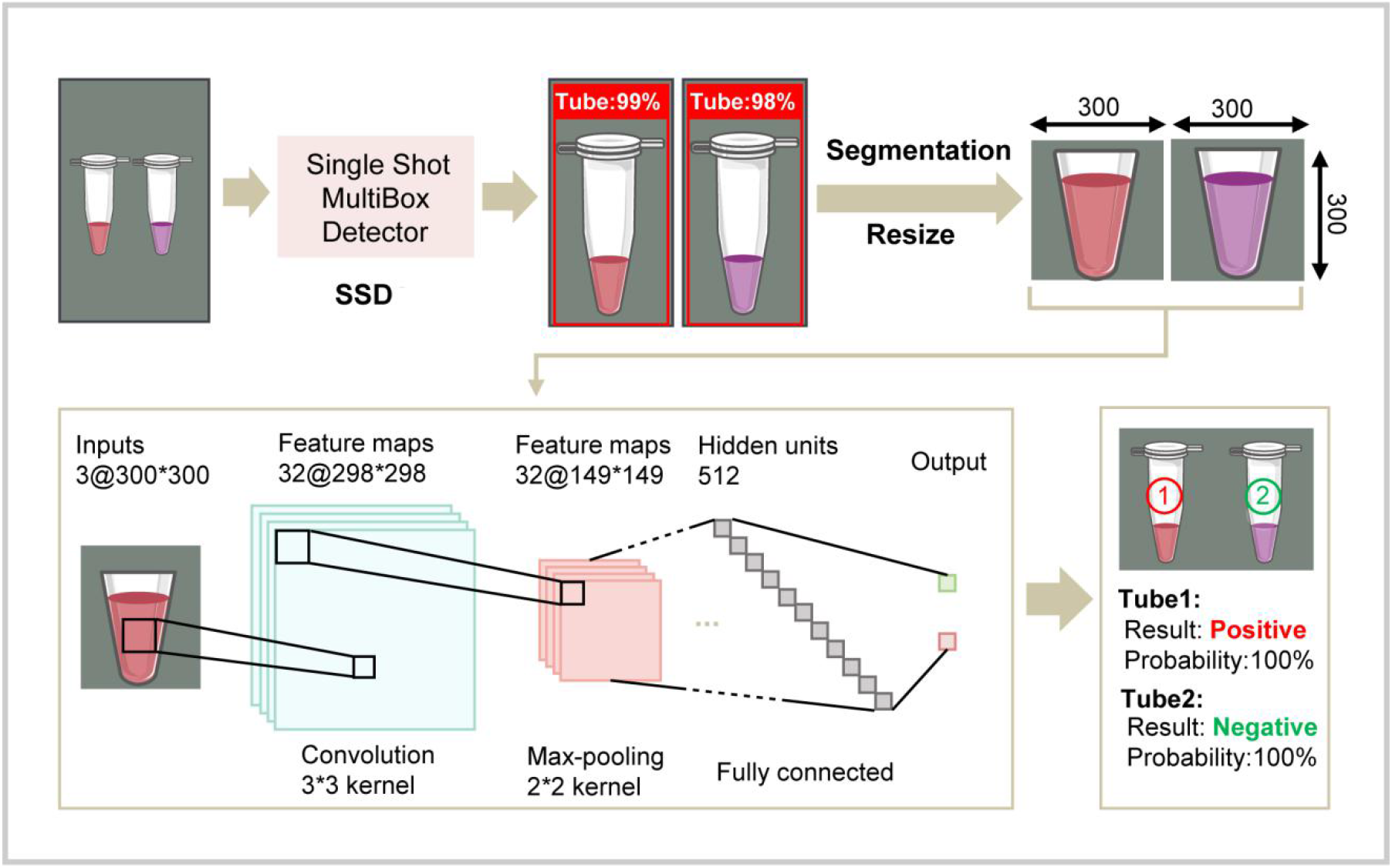
Schematic diagram of analysis of the RAVI-CRISPR assay read-outs by two convolutional neural network-based machine learning models. Single Shot MultiBox Detector (SSD) algorithm identification of reaction tubes and binary classification of positive or negative detection signals.

To train the first model to recognize the images of PCR tubes, we used 1554 single-PCR tube images, 405 multiple PCR tube images, 283 8-tube PCR strip images, and 301 mixed images (8-tube PCR strip mixed with single-PCR tube images) which were manually labeled (Supplementary Table S4). Testing of the object detection model showed that this algorithm could identify PCR tubes in all cases (Supplementary Fig. S4) with the detection rates at 100% for both single and multiple PCR tube images and > 99% for 8-tube PCR strip images (Supplementary Table S5). Subsequently, the binary classification model was trained to distinguish colorimetric changes in the tubes containing the RAVI-CRISPR reactions with either positive or negative detection signals (Supplementary Table S6). We found that the F1-Measure, which was used to evaluate the accuracy of predictions in two-class (binary) classification problems, could be increased to 100% by using an optimal ROX-labeled reporter in concentration > 1 μM (Supplementary Fig. S5). This result meant that the model performed well on images captured with no excitation light when the probe concentration was relatively high (Supplementary Fig. S6).

In order to evaluate the robustness of the model at critical concentrations of DNA samples, using images captured under no excitation light as an example, we tested the model on samples with probe concentrations of 1 μM and DNA concentrations ranging from 0 to 7 × 10^5^ copies/μL. The result showed that even at very low concentration of DNA template, the model could discriminate negative and positive signals with 100% accuracy (Supplementary Fig. S7, Table S7). We also found that the sensitivity of the image analysis app was much higher than that of the naked eye read-out, indicating the power of the app to be used in circumstances where the naked eye evaluation is the only option.

To further evaluate the classification model, we examined the ConvNet outputs at the intermediate training stage of the convolutional model and found that many activation channels were focused on the tip of the PCR tube (Supplementary Fig. S8), suggesting that the model may classify results primarily by distinguishing tip color where the signal was the strongest, in the same way as the naked eye observation. Finally, we deployed this algorithm on a cloud server as the MagicEye mobile application for accurate and rapid analysis of nucleic acid detection signals.

### Single-tube RAVI-CRISPR assay and mobile MagicEye system for POCT of nucleic acids

The two-step RAVI-CRISPR assay requires opening of the tube after RT-LAMP or LAMP reaction, which may generate aerosols, potentially causing false-positive results. To reduce the likelihood of contamination and improve the convenience of visualization process for POCT of COVID-19 or ASF, we next established a single-tube RAVI-CRISPR assay. The underlying principle and operation of the assay relied on an isothermal amplification with CRISPR/Cas12a-based detection (Fig. 5A). To avoid cross-contamination, mineral oil was added to cover the extracted nucleic acids and isothermal amplification solution. Cas12a and crRNA ribonucleoproteins (RNPs) reagents were pre-added inside of the tube lid. Following the isothermal amplification, RNPs reagents were mixed with amplification reagents by hand shaking (Fig. 5A). The whole process did not require the lid to be re-opened, thus avoiding the possibility of aerosol contamination.

**Figure 5.**
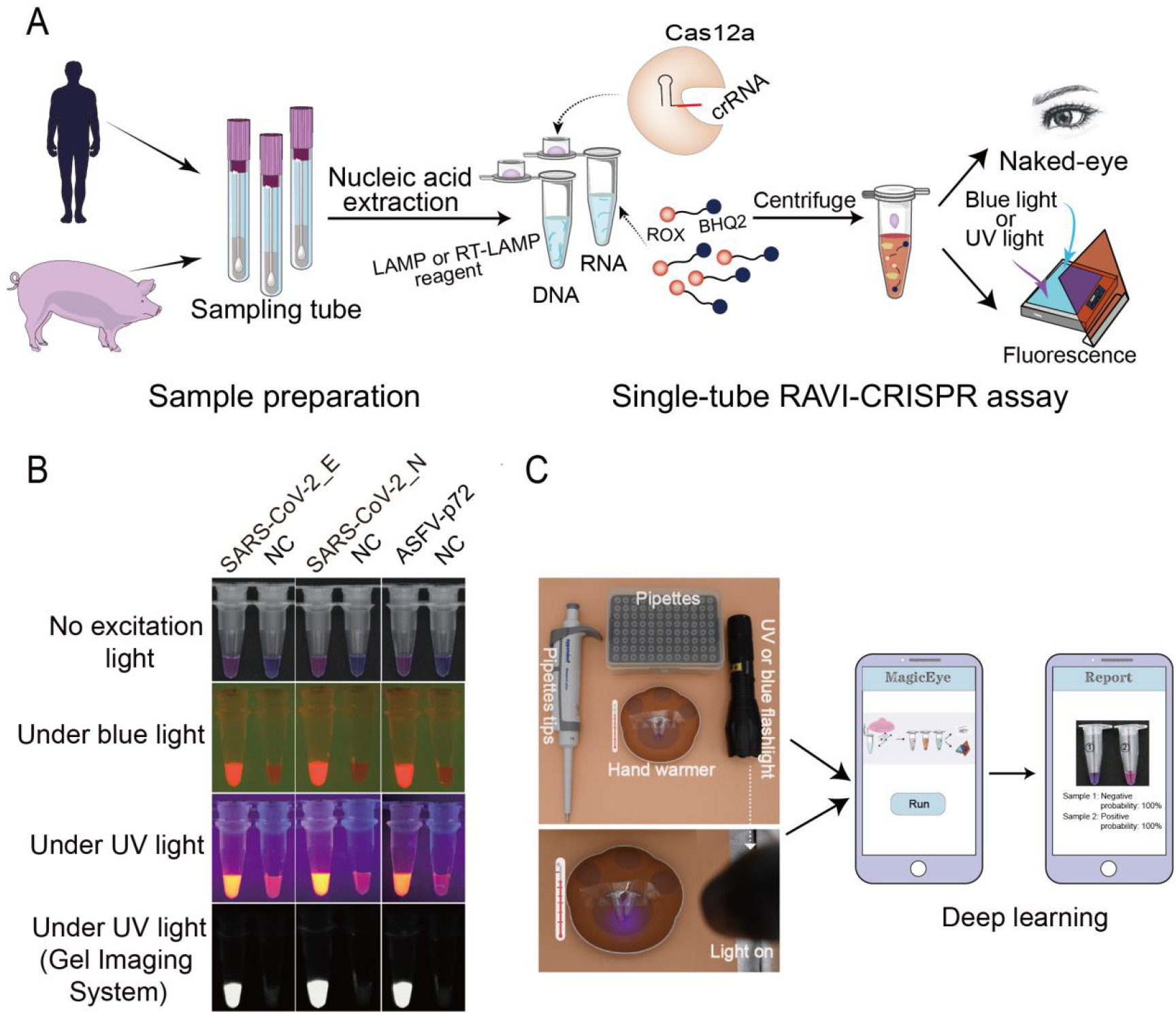
Establishment of the contamination-free single-tube RAVI-CRISPR assay and mobile MagicEye system for a point-of-care detection of nucleic acids. **A.** Diagram of overall experimental flow and principle of the contamination-free single tube RAVI-CRISPR assay. **B.** Visual detection of the SARS-CoV-2 *E* and *N* genes as well as the ASFV *p72* gene by the single-tube RAVI-CRISPR assay. **C.** Photo display of portable rechargeable hand warmers and smart phones for point-of-care testing.

We next tested the combined single-tube RAVI-CRISPR/MagicEye system with simulated *in vitro* transcripts or real clinical samples and found that the SARS-CoV-2 *E* and *N* genes and the ASFV *p72* genes were all successfully detected (Fig. 5B). To facilitate the pen-side, instrument-less detection of virus, we used a portable rechargeable hand warmer to provide temperatures from 35 to 60 ℃ for the amplification incubation. To enhance the fluorescence signal for the pen-side detection, a small flashlight capable of UV and blue light emission were approved to be sufficient (Fig. 5C). Analysis by a smartphone with the MagicEye app successfully replicated the result of detection by the naked eye for these samples (Supplementary Fig. S9, Table S8), thereby demonstrating this single-tube RAVI-CRISPR assay instrumented only with portable rechargeable hand warmer and MagicEye app to be readily deployed in the pen-side POCT of for SARS-CoV-2 or ASFV among other viral pathogens.

## Discussion

Current and future global pandemics of viral infections, like the ongoing COVID-19 pandemic caused by SARS-CoV-2, pose overwhelming threats to human health and global economic stability. Thus, the development of affordable and convenient POCT diagnostic tools is both warranted and urgent. In the present study, we have developed a rapid, single-tube, instrument-less, colorimetric POCT assay (“RAVI-CRISPR”) for the detection of nucleic acids as a demonstration of the proof-of-concept for its sensitive detection of diverse pathogenic nucleic acids by the naked eye. We also developed the MagicEye mobile app for standardized diagnostic image analysis of color changes in the reaction tubes. The main feature of this POCT assay is that ssDNA-FQ reporters are used as colorimetric sensors to produce changes in the color of the reaction mixture dependent on the concentrations of the reporters and specifically targeted nucleic acids that are cleaved by Cas12a. To our knowledge, this is the simplest CRISPR based-nucleic acid platform reported to date, since no transducing component is required for visualization (Supplementary Table S9).

Although previous studies have reported that LAMP reaction can amplify targeted nucleic acids with high efficiency, this reaction produces long concatemeric amplicons, which make sequence-specific detection of LAMP products analytically challenging^26^. A recent study reported that the use of CRISPR/Cas12a in combination with a sequence-specific plasmonic LAMP assay can provide orthogonal color read-outs^47^. However, plasmonic gold nanoparticles (AuNPs) are difficult to produce and their waste is hard to manage, thus limiting their application. To resolve these issues, ssDNA-FQ reporters are proved to be straightforward to synthesize and environmentally harmless compared to AuNPs. We found that the ssDNA-FQ reporter concentration exerts a strong impact on the direct visual detection of nucleic acids. By adjusting the concentration of ROX-labeled reporter, a clear naked-eye observation of the detected nucleic acids became feasible without excitation light. In comparison, colorimetric detection by our RAVI-CRISPR system using ROX-labeled reporter produced stronger color changes than that of the AIOD-CRISPR assay, which used FAM-labeled reporters^44^. In particular, our optimization process identified eight novel ssDNA-FQ reporters suitable for direct assessment with no excitation light.

To facilitate the standardization for colorimetric image detection, we also developed a cloud server-based, convolutional neural network algorithm (MagicEye app) for automatic analysis of the testing results. Although two apps, HandLens^41^ and STOPCovid.v2^42^, were previously developed for CRISPR-based nucleic acid detection, these apps only focused on image analysis of lateral flow strip read-outs, whereas MagicEye can recognize and analyze colorimetric changes of single or 8-strip PCR tubes in multiple test reactions (Supplementary Table S9). Our MagicEye can also improve the detection sensitivity of RAVI-CRISPR assays. The accurate diagnosis of ASFV infection in pigs by MagicEye and visual inspection confirmed the reliability of the RAVI-CRISPR assay. This assay was successfully used for the single-copy detection of ASFV samples in a single-tube incubated only by a portable rechargeable hand warmer. Moreover, this system does not require an electrical power source, therefore it can be feasibly deployed for pen-side testing in rural areas with limited resources.

## Materials and Methods

### Nucleic acid preparation

The partial fragment 780 bp of ASFV (GenBank accession number: MK 333180) *p72* gene was synthesized (TSINGKE, Beijing, China) and cloned into the pMD18T vector (Cat. No. D101A, TAKARA, China). pBluescript-N vector containing 931 bp of SARS-CoV-2 (GenBank accession number: NC_045512.2) *N* gene was acquired from Fubio (Cat. No. FNV2614, Shanghai, China). The *E* gene 374 bp of SARS-CoV-2 was synthesized and cloned into the pUC57 vector (TSINGKE, Beijing, China). The *N* gene 235 bp and *E* gene 563 bp of SARS-CoV (GenBank accession number: NC_004718.3) were synthesized (TSINGKE, Beijing, China). The *N* gene 232 bp and *E* gene 249 bp of MERS-CoV (GenBank accession number: KT326819.1) were also synthesized (TSINGKE, Beijing, China) (Supplementary Table S10).

For ASFV, blood samples were collected from pigs before and at different time points of an experimental challenge by ASFV at the International Livestock Research Institute (ILRI), Kenya. DNA was extracted from the EDTA-treated blood samples using the DNeasy Blood & Tissue Kits (Cat. No. 69504, Qiagen, GmBH, Germany) at the BSL2 lab of ILRI. Both qPCR negative and qPCR positive samples were selected to test the system.

### Exonuclease I cleavage assay

Sixteen ssDNA-FQ reporters, namely, ROX-N12-BHQ2, Texas Red-N12-BHQ2, VIC-N12-BHQ2, JOE-N12-BHQ1, HEX-N12-BHQ1, TAMRA-N12-BHQ2, TET-N12-BHQ1, PET-N12-BHQ1, Alexa Flour 633(NHS)-N12-BHQ2, Quasar 570-N12-BHQ2, Quasar 670-N12-BHQ3, FAM-N12-BHQ1, Cy3-N12-BHQ2, Cy5-N12-BHQ2, Cy5.5-N12-BHQ2, and Cy7-N12-BHQ2 were synthesized (TSINGKE, Beijing, China) for the optimization of their concentration range and gradient in CRISPR-Cas12a reaction. All these ssDNA-FQ reporters contained a fluorophore at their 5′ end and a matched non-fluorescent quencher at their 3′ end. The ssDNA-FQ reporters are listed in Supplementary Table S10. Exonuclease I (Cat. No. M0293S, New England Biolabs (NEB), Ipswich, MA, USA) cleavage was performed for different concentrations of ssDNA-FQ reporters at 37 °C for 15 min on the NaCha Multi-Temp Platform (NaCha, Monad, Suzhou, China). The signals of degraded ssDNA-FQ reporters after the cleavage were detected by the visible light gel viewer (EP2020, BioTeke, Wuxi, China) and the automatic gel imaging analysis system (Peiqing Science and Technology, Shanghai, China) with blue and UV light, respectively. The fluorescence intensity and background signals of different concentrations of the ssDNA-FQ reporters before and after the cleavage were further detected by the EnSpire Multimode Plate Reader (EnSpire, PerkinElmer, Waltham, MA, USA). Fluorescence excitation and emission wavelengths were set to 490–750 and 520– 770 nm, respectively, which corresponded to the excitation and emission profiles of the ssDNA-FQ reporters, respectively.

### Design of target sites for virus-specific crRNAs

Five crRNAs targeting SARS-CoV-2 *E* and nine crRNAs targeting SARS-CoV-2 *N* genes were designed by CRISPR-offinder software^48^ (Supplementary Table S10) and complemented to the target sites with a 5ʹ “TTTN” PAM in the DNA strand opposite the target sequence (Supplementary Table S10). A highly active crRNA against ASFV *p72* gene was selected from our previous study37.

### *In vitro* RNA transcription using T7 RNA polymerase

SARS-CoV-2 *N* and *E* genes were transcribed from the pBluescript-N and pUC57-E plasmids by adding a T7 promoter via PCR using Premix Taq (Cat. No. R004A, TAKARA, Shuzo, Shiga, Japan). The crRNA templates were amplified from a pUC57-T7-crRNA (Supplementary Table S10) using a forward primer containing a T7 promoter and a reverse primer containing the nucleotide target sequences in different lengths (Supplementary Table S10). The PCR products were purified using the PCR purification kit (Cat. No. TD407, Beijing Tianmo Sci & Tech Development, Beijing, China) and then served as the DNA template for *in vitro* transcription reactions using the HiScribe T7 High Yield RNA Synthesis Kit (Cat. No. E2040S, NEB). Template DNA was removed by the addition of DNase I (Cat. No. M0303S, NEB) and *in vitro* synthesized RNA was subsequently purified using Monarch RNA Cleanup Kit (Cat. No. T2040S, NEB). RNA concentration was quantified by NanoDrop 2000 spectrophotometer (Thermo Fisher Scientific, Wilmington, DE, USA) and copy number was calculated using their transcript length and concentration.

### RT-qPCR

Different copies of RNAs transcribed *in vitro* from SARS-CoV-2 *N* and *E* genes were quantified via RT-qPCR. The primers, probes, and their relative concentrations were those recommended by the Centers for Disease Control and Prevention (CDC) and WHO (Supplementary Table S10). The Luna Universal Probe One-Step RT-qPCR Kit (Cat. No. E3006S, NEB) was used as the relevant master mix, RT-qPCR reactions were set up according to the manufacturer’s instructions, and the thermocycling settings were according to the CDC and WHO protocol^49,50^. *N* and *E* gene standards were used to generate a standard curve for copy number quantification. The qPCR assay and primers (Supplementary Table S10) for the detection of ASFV *p72* followed the reaction conditions recommended by OIE^51^. SYBR Green Realtime PCR Master Mix (Cat. No. QPK-201, Toyobo, Osaka, Japan) was used as the relevant master mix. The qPCR reaction was performed in the CFX96 Touch Real-Time PCR Detection System (Bio-Rad, Hercules, CA, USA).

### RAVI-CRISPR assays with RT-LAMP/LAMP

RT-LAMP/LAMP primers (Supplementary Table S10) used the amplification of the fragments 238 bp and 200 bp of both *N* and *E* genes of SARS-CoV-2 as well as the *p72* gene 198 bp of ASFV were designed using PrimerExplorer V5 (https://primerexplorer.jp/e/). The reaction was performed on a series of dilutions, and the products were analyzed by gel electrophoresis. Each amplification template containing appropriate copy number of the gene was added into the master mix containing 1 μL of Bst 3.0 DNA polymerase (Cat. No. M0374L, NEB), 2.5 μL of 10 × isothermal amplification buffer, 6 mM MgSO4, 14 mM each of dNTP Mix, and 2.5 μL of primer mix (1.6 μM FIP/BIP, 0.2 μM F3/B3, and 0.4 μM LF/LB). The amplification reactions were performed in the NaCha Multi-Temp Platform at 65 °C for 40 min. Then, 3 μL of each sample was used for 2% agarose gel electrophoresis and 3 μL of the remaining portion was set aside for RAVI-CRISPR assays.

EnGen^®^ Lba Cas12a (Cat. No. M0653S, NEB) and NEBuffer 2.1 (Cat. No. B7202S, NEB) were purchased from the NEB. Each sample of RAVI-CRISPR assay contained 500 nM Cas12a, 1 μM crRNA, 20 μM ssDNA-FQ reporter (ROX-N12-BHQ2), 3 μL of pre-amplificated product, and 2 μL of NEBuffer 2.1. The total volume was adjusted to 20 μL with nuclease-free water. The reactions were incubated at 37 °C for 15 min and inactivated at 98 °C for 2 min. The reverse primers of *in vitro* transcription containing crRNA complementary sequences were used as ssDNA activators (Supplementary Table S10). PCR tubes containing 20 μL of the RAVI-CRISPR assay samples were placed on a black board (under no excitation light) or under a blue/UV light transilluminator for the naked eye or fluorescence visualization. The results were recorded using a smartphone camera or the Peiqing automatic gel imaging analysis system with the same exposure time. For fluorescence intensity measurement by the EnSpire Multimode Plate Reader, 2.5 μM ssDNA-FQ reporters were added to the tubes, and fluorescence excitation and emission wavelengths were set to 576 and 601 nm, respectively. For lateral flow visual detection, 2.5 μM ssDNA-FQ (FAM-N12-Biotin) reporter was added to the tube, followed by the treatment with the HybriDetect - Universal Lateral Flow Assay Kit (Milenia Biotec, Gießen, Germany), and the result was visualized after approximately 5 min.

### Single-tube RAVI-CRISPR assay

Single-tube RAVI-CRISPR assay was conducted using components A and B. Component A contained 1 μL of Bst 3.0 DNA polymerase (Cat. No. M0374L, NEB), 2.5 μL of 10 × isothermal amplification buffer, 6 mM MgSO4, 14 mM each of dNTP Mix, and 2.5 μL of primer mix (1.6 μM FIP/BIP, 0.2 μM F3/B3, and 0.4 μM LF/LB). Component B contained 500 nM Cas12a, 1 μM crRNA, and 2 μL of NEBuffer 2.1. For the 25 μL of single-tube RAVI-CRISPR assay, component A was first added to the bottom of the PCR tube. Then, the amplified template mixed with 20 μM ssDNA-FQ (ROX-N12-BHQ2) reporter was added to the PCR tube and vortexed to mix. Next, 1.6 μL of mineral oil was added to the bottom liquid for insulation, then 5 μL of component B was placed in the lid, and PCR was carried for 40 min at 65 °C. Afterward, the PCR tube was mixed thoroughly and reacted at 37 °C for 15 min. All incubation reactions were performed on the portable rechargeable hand warmer, which provided temperatures from 35 to 60 °C. For the naked eye or fluorescence visualization, PCR tubes containing 25 μL of single-tube RAVI-CRISPR reaction solution were checked under blue or UV light.

### Image identification and segmentation in PCR tubes

A single deep neural network, SSD (https://github.com/bubbliiiing/ssd-keras)^46^, was used to identify the images of PCR tubes, and the detection models were trained based on the Keras (v2.1.5) and TensorFlow (v1.13.1) frameworks^52,53^. The images captured under no excitation light were used to train the PCR tube recognition models. Single-PCR tube and 8-tube PCR strip were used to improve the generalization power of the models. In addition, functions in the opencv-python (v4.4.0.46) library (https://github.com/opencv/opencv-python), namely, “median blur operation”, “opening operation”, “closing operation”, “binary thresholding operation”, “kmeans”, “PCACompute2”, and “findContours”, were used to locate and segment the images of the PCR tube tip to maximize the utilization of effective information of the images under no excitation light.

### Construction of a binary classification model for positive and negative detection signals

A convolutional neural network (CNN)-based model was constructed for the binary classification of positive and negative detection signals. These CNN models were trained based on the Keras (v2.1.5) and TensorFlow (v1.13.1) frameworks. Then, we referred to the suggestions of François Chollet in “Deep Learning with Python” to regularize the model and optimize the hyperparameters as our previous research^54,55^. The CNN model consisted of four convolutional layers with 2D convnets, four pooling layers, and two fully connected layers. The “filter” parameters for convolutional layers were 32, 64, 128, and 128, and the remaining parameters were set as “kernel_size = (3,3), activation = relu”. The parameters for pooling layers were set as “pool_size = (2,2)”. For fully connected layers, after 512-way rectified linear unit (ReLU) layers, a one-way sigmoid layer was designed to return a probability score. Additionally, we exposed the model to multiple aspects of the images through data augmentation, such as image rotation, image shift, and image shear, to enhance the generalization power. The parameters were: “rescale = 1./255, rotation_range = 40, width_shift_range = 0.2, height_shift_range = 0.2, shear_range = 0.2, zoom_range = 0.2, horizontal_flip = True”. In addition, MagicEye mobile phone app (Android version 1.0) was developed and can be freely downloaded from “https://sourceforge.net/projects/ravi-crispr/files/MagicEye_en_v1.0.apk/download”.

### Statistical analysis

Statistical analysis was performed using R programming language. Mean ± standard errors of the mean (SEM) was determined for each treatment group in the separated experiments. The two-tailed Student’s t test was used to determine significant differences between treatment and control groups (^***^*P* < 0.001, ns: no significant at the 95% confidence level).

## Data Availability

All study data are included in the article and Supplemental Information.

## Acknowledgements

This work was supported by the NSFC Major Research Plan-Major Scientific Problems of African Swine Fever virus (31941008), the Natural Science Foundation of China (32072685), and the China Agriculture Research System of MOF and MARA. We thank Dr. Xiangru Wang for technical support. We also thank the CGIAR Research Program on Livestock and the CGIAR Consortium for support.

## Author Contributions

Most of the experimental work and data analysis were co-conducted by D.T., B.X., and S.X., with minor contributions from W.P., X.H., C.Z., J.R., L.F., X.N., Y.Z., Y.M., and J.H. Y.F. developed convolutional neural network assessment of RAVI-CRISPR assay read-outs. Y.T. developed the MagicEye mobile application. S.X. conceived the project, designed the experiments, and wrote the manuscript. S.Z., X.L., X.L, and S.L. provided support and supervised the project. All authors contributed to manuscript revision.

